# Cannabidiol (CBD) potentiates physiological and behavioral markers of hypothalamic-pituitary-adrenal (HPA) axis responsivity in female and male mice

**DOI:** 10.1101/2024.10.17.618869

**Authors:** Bryan W. Jenkins, Hayley A. Spina, Kate Nicholson, Amy E. M. Newman, Jibran Y. Khokhar

**Affiliations:** Division of Behavioral Biology, Department of Psychiatry and Behavioral Sciences, Johns Hopkins University School of Medicine; Department of Integrative Biology, College of Biological Sciences, University of Guelph; Department of Biomedical Sciences, Ontario Veterinary College, University of Guelph; Department of Anatomy and Cell Biology, Schulich School of Medicine and Dentistry, Western University Canada

**Author notes:** Corresponding author: Jibran Y. Khokhar, Ph. D., Associate Professor, Canada Research Chair in Translational Neuropsychopharmacology, Anatomy and Cell Biology, Schulich School of Medicine and Dentistry, Western University, Faculty, Graduate Program in Neuroscience. Authors contributed equally (Bryan W. Jenkins, Hayley A. Spina). Funding: Funding was provided by the University of Guelph (AEMN) as well as the Natural Sciences and Engineering Research Council (RGPIN-19-05121; JYK).

**Keywords:** phytocannabinoid, stress, cognition, affective disorders, sex differences, anxiety

## Abstract

**Rationale:** Clinical literature indicates there may be a therapeutic use of cannabidiol (CBD) for stress-related disorders. Preclinical literature remains conflicted regarding the underlying neurobehavioral mechanisms, reporting mixed effects of CBD (increased, decreased, or no effect) on anxiety- and fear-related behaviors. Preclinical data demonstrated that CBD modulates hypothalamus-pituitary-adrenal (HPA) axis gene expression; it is unknown whether CBD changes HPA axis responsivity and how this relates to altered behavior.

**Objectives:** We aimed to evaluate whether acute or chronic CBD administration would alter physiological and behavioral measures of HPA axis responsivity in male or female mice.

**Methods:** C57BL/6 mice of both sexes were injected with vehicle or CBD (30 mg/kg, i.p.) daily for 26 days. Plasma corticosterone (CORT) levels were evaluated following dexamethasone suppression and adrenocorticotropin hormone stimulation tests after acute and chronic CBD exposure. After chronic CBD, mice were tested for anxiety-like behavior using an elevated plus maze (EPM) and associative fear learning and memory using a trace fear conditioning (FC) protocol.

**Results:** Compared to vehicle, CBD induced a state of HPA axis hyperactivation, an effect which was significant in males; it also normalized anxiety-like behavior in female mice classified as having HPA axis hypofunction and primed all female mice for enhanced conditioned responding. Significant sex differences were also detected: females had greater plasma CORT levels and HPA axis responsivity than males, exhibited less EPM anxiety-like behavior, and were more responsive during FC.

**Conclusions:** CBD potentiated physiological and behavioral markers of HPA axis function and normalized anxiety-like behavior in a sex-specific manner. This observation has implications for cannabinoid-based drug development targeting individuals with stress-related disorders involving HPA axis hypofunction pathology.

## INTRODUCTION

Mental health disorders involving maladaptive stress and fear responses are highly prevalent in the United States. According to Harvard’s National Comorbidity Survey, nearly 20% of adults had an anxiety disorder and nearly 4% had post-traumatic stress disorder (PTSD) in 2007 (Harvard Medical School, 2007). Prevalence rates for anxiety-related disorders and PTSD are also greater for females than males (Kessler et al., 2005; McLean et al., 2011). Patients with anxiety disorders or PTSD may experience adverse responses to medication or may not respond at all (Ballenger, 2004; Krystal et al., 2011; Otto et al., 2001). Thus, novel pharmacotherapeutics are urgently needed to improve outcomes for patients (Blessing et al., 2015).

Cannabidiol (CBD) is the second most studied phytocannabinoid in *Cannabis sativa* and may have therapeutic potential, including for the alleviation of anxiety and PTSD symptoms (J. A. Crippa et al., 2018; Guimarães et al., 1990; Papagianni & Stevenson, 2019; Uhernik et al., 2018). In healthy humans and in patients with generalized social anxiety disorder, oral CBD reduced self-report measures of anxiety at baseline and during the Simulated Public Speaking Test. (Bergamaschi et al., 2011; J. A. Crippa et al., 2004; J. A. S. Crippa et al., 2011; A. W. Zuardi et al., 1993). Similarly, patients being treated for PTSD had improved outcomes after eight weeks of adjunctive oral CBD (Elms et al., 2019). Clinical trials are now investigating CBD pharmacotherapies (García-Gutiérrez et al., 2020).

Preclinical studies have used rodent models to evaluate the behavioral and neurobiological mechanisms of CBD effects on anxiety-like behavior and associative fear learning, with mixed results. In female and male mice, CBD has either dose-relatedly increased, decreased, or had no effect on anxiety-like behavior (Huffstetler et al., 2023; Kasten et al., 2019; Liu et al., 2022; Sturaro et al., 2023; Zieba et al., 2019). Similarly, CBD has enhanced, attenuated, or had no effect on associative fear learning and memory (Assareh et al., 2020; Chesworth et al., 2022; Montoya et al., 2020; Uhernik et al., 2018). Investigating changes in underlying stress-related neurobiology may reveal novel mechanisms and help resolve the conflicting preclinical data.

A logical target for such mechanistic investigations is the hypothalamic-pituitary-adrenal (HPA) axis. The HPA axis is the primary neuroendocrine pathway that regulates the physiological response to a stressor through the release of glucocorticoids (cortisol in humans and corticosterone [CORT] in rodents). At baseline, daily rhythmicity of CORT functions to regulate energy availability (Dallman et al., 1993) and cognition including learning and memory (Liston et al., 2013; Rimmele et al., 2010). Following exposure to a stressor, increases in CORT functions to mobilize several physiological systems which maintain homeostasis and facilitate recovery following removal of the stressor (Sapolsky et al., 2000). Chronic stress can alter HPA axis functioning, resulting in changes to both baseline and stress-induced CORT release (as reviewed by (Herman et al., 2016)). Dysregulated HPA axis functioning is associated with some anxiety-related disorders and PTSD, though the mechanisms for how altered HPA axis regulation contributes to the pathology of these disorders remains debated (de Kloet et al., 2006; Plag et al., 2013).

Recently, Viudez-Martínez et al. (2018) demonstrated that CBD treatment blocked restraint stress-induced changes in corticotropin-releasing factor and glucocorticoid receptor gene expression in the paraventricular nucleus and hippocampus of mice (Viudez-Martínez et al., 2018). Because this study demonstrated the potential for CBD to regulate HPA axis gene expression, we hypothesized that CBD administration would alter physiological and behavioral markers of HPA axis activation. We specifically hypothesized that effects of CBD administration would be evident after acute exposure and would decrease with chronic exposure as HPA axis signaling adapts to on-board CBD. We further hypothesized that effects of CBD on HPA axis responsivity and behavior would be sex-dependent. This work is important as it will identify a potential mechanism of action and sex-dependent effects of CBD relevant for treating stress-related disorders.

## METHODS

### Animals

Thirty-two male and female adult C57BL/6J mice (n = 16 for each sex) were obtained from Charles River Laboratories (Senneville, Quebec, Canada). Sex- and treatment-matched mice were housed in polyethylene cages (n = 4 per cage; Allentown, Allentown, New Jersey, United States) with environmental enrichment and *ad libidum* access to food and water. Mice were maintained on a 12:12 hour light:dark cycle (on/off at 7:00 AM/PM) in a room with circulating air, a constant ambient temperature of 22 ± 2 ^o^C, and humidity of 50-70%. All experiments were approved by the University of Guelph Institutional Animal Care and Use Committee and performed in accordance with the guidelines described in the Guide to the Care and Use of Experimental Animals (Canadian Council on Animal Care, 1993).

### Drugs

Cannabidiol (CBD; Toronto Research Chemicals, Toronto, Ontario, Canada) was dissolved in 100% ethanol, tween-80, and saline (0.9% NaCl) in a 1:1:18 ratio. The vehicle contained the same amounts of ethanol, tween-80, and saline without the added CBD. Drugs were prepared fresh every two or three days. Dexamethasone (DEX; Sigma-Aldrich, Oakville, Ontario, Canada) and adrenocorticotropic hormone (ACTH; Abcam, Waltham, Boston, USA) were each dissolved in a mixture of 100% ethanol and saline (1:19 ratio). Mice were weighed daily before injections. Mice received daily intraperitoneal (i.p.) injections of either vehicle or CBD (30 mg/kg) at a volume of 1 ml/kg starting at 8:00 am (n = 8 per sex and treatment) for 26 days. This dose was selected based on data demonstrating altered anxiety-like behavior and fear conditioning in mice (Assareh et al., 2020; Fogaça et al., 2018) and modest physiological relevance for therapeutic efficacy in humans based on preclinical pharmacokinetic data (Kwee et al., 2022).

### Dexamethasone Suppression Test, ACTH Stimulation Test, and Sample Collection

Mice underwent DEX suppression followed by ACTH stimulation to assess HPA axis responsivity at acute (Day 1) and chronic (Day 14) timepoints (Fig 1A). One hour after vehicle or CBD injections, mice were given injections of DEX (0.1 mg/kg, i.p.). Mice were placed back in their home cages and left undisturbed for four hours, after which a blood sample was collected from the saphenous vein. Immediately following blood collection, mice were administered ACTH injections (0.5 mg/kg, i.p.) and placed back in their home cages. After one hour, blood was again collected from each mouse via the saphenous vein. Following the end of behavioral testing, mice were sacrificed via cervical dislocation and trunk blood was collected to measure plasma CORT levels. All blood samples were stored on ice for approximately one to two hours, then centrifuged at 12,000 rpm for 10 min and serum was collected and stored at −20 °C until subsequent analysis.

**Fig. 1.**
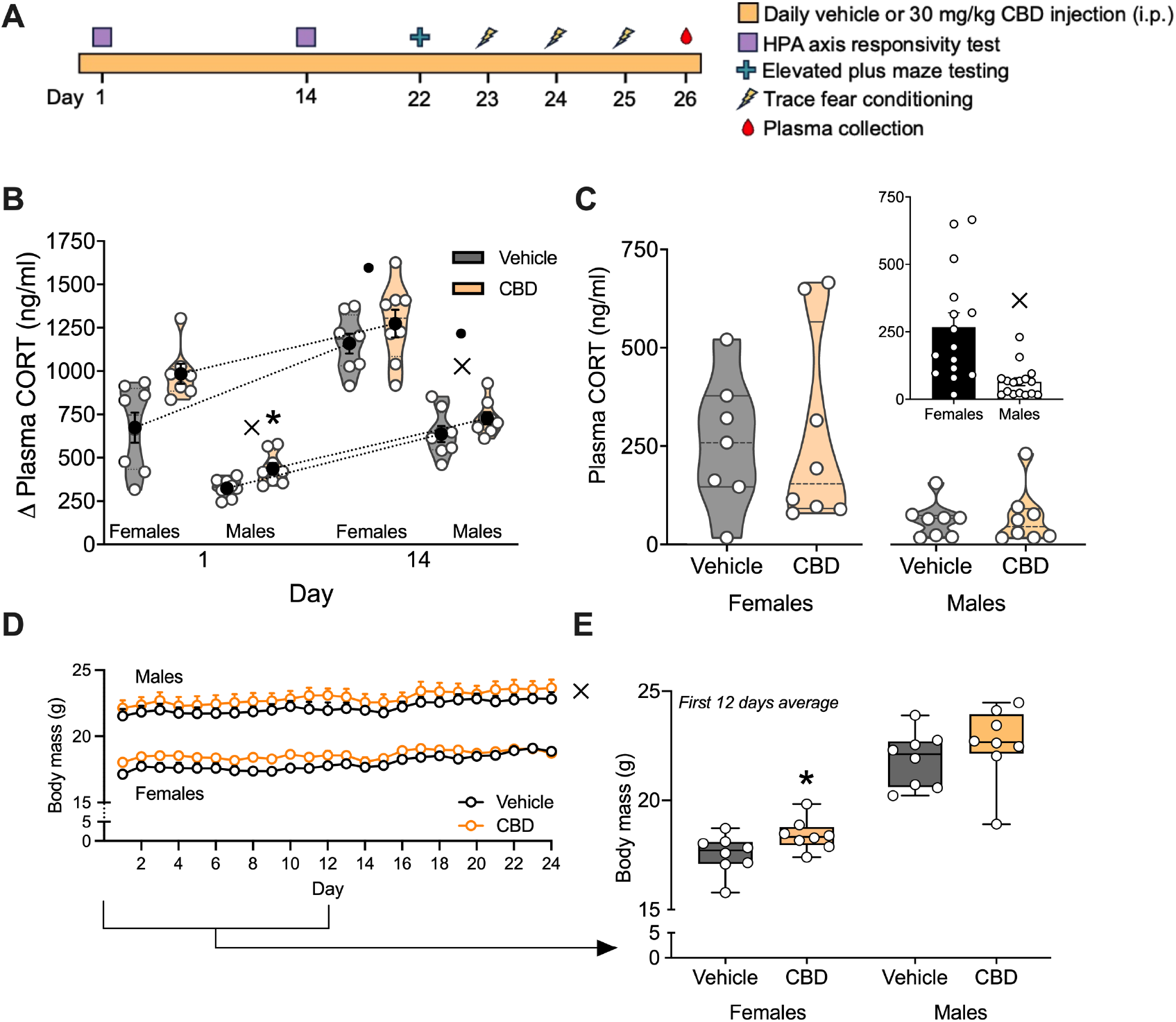
Experimental timeline and CBD effects on HPA axis physiology. **A)** Experimental timeline: mice (n = 8/sex/group) underwent daily vehicle or CBD (30 mg/kg, i.p.) injections for 26 days (orange rectangle). HPA axis responsivity testing occurred at acute (day 1) and chronic (day 14) timepoints (purple squares); EPM (blue plus) and trace FC (yellow lightning bolts) testing was between Days 22-25, after which plasma samples were collected (red droplet). **B)** CBD (orange violin plots) increased HPA axis responsivity (△ Plasma CORT) compared to vehicle (grey violin plots), which was significant in males and resolved by day 14; black circles with error bars represent mean ± SEM. HPA axis responsivity was greater in females and at the chronic timepoint. **C)** Chronic CBD did not affect plasma CORT levels (violin plots); female CORT levels were greater than males regardless of CBD exposure (**C inset panel** = data collapsed across CBD condition). **D)** Compared to vehicle (grey time series), daily CBD (orange time series) increased body mass in females only, which resolved by day 14; males overall weighed more than females. **E)** Daily CBD (orange boxplot) increased average body mass over the first 12 days of testing compared to vehicle (grey boxplot) in females only. HPA = hypothalamus-pituitary-adrenal, EPM = elevated plus maze, FC = fear conditioning, CORT = corticosterone, CBD = cannabidiol, * = compared to vehicle, × = compared to female, • = compared to earlier timepoint.

### Elevated Plus Maze

Mice were assessed for anxiety-like behaviors using the elevated plus maze after 21 days of chronic vehicle or CBD treatment (Fig 1A). Mice were placed in the center of the maze facing one of two open arms and had five minutes to explore the two open and two closed arms of the maze. The amount of time spent exploring and the distance travelled in each of the four arms were tracked in 60-second time bins using Ethovision XT 16 (Noldus Information Company, Leesburg, Virginia, USA). The number of entries to the open arms of the maze, the speed in each arm, and the latency to enter the open arm were also computed. Stretch-attend postures were detected for each animal using an elongation threshold of 80% and summed.

### Trace Fear Conditioning

Mice were tested for associative fear learning and memory using a trace fear conditioning protocol adapted from Curzon et al., (2009). At least 24 hours prior to testing, mice were habituated to the experiment room for 60 minutes. On each test day (Fig 1A), mice were placed in four fear conditioning chambers (Med Associates Inc., St. Albans, Vermont, USA) housed within sound-attenuating cabinets, with one of two types of electrified floor bars to deliver a shock as the unconditioned stimulus (0.6 mA, US). Speakers located beside the fear conditioning chambers presented background noise (67 dB white noise) and an auditory cue as the conditioned stimulus (90 dB pure tone, CS). Each fear conditioning chamber was outfitted with different odors (vanilla or lemon extract) and spatial geometries (standard or curved walls and parallel or zig-zag floor bars) as contextual cues. Cues were counterbalanced across animals in each experiment group.

The first test day consisted of a conditioning day whereby mice were exposed to three pairings of the CS (i.e., the auditory tone) with the US (i.e., the foot shock) to initiate associative fear learning. The second test day consisted of a context testing day whereby mice were exposed to the same contextual cues in which they were conditioned, without the presentation of the paired CS + US, to evaluate contextual fear memory recall. The third test day consisted of a cue testing day whereby mice were exposed to the CS in an environment with novel contextual cues, to evaluate cued fear memory recall. Freezing behavior was recorded and analyzed using the accompanying Video Freeze (Med Associates Inc.) software. Data were collected continuously and segmented into a 180-second baseline and three 60-second post-CS/US time bins for conditioning testing, eight 60-second time bins for context testing, and a 30-second baseline and ten 60-second post-CS time bins for cue testing (sampled every 30 seconds).

### Corticosterone Assays

CORT levels in serum were measured using an I125 Corticosterone Double Antibody RIA Kit (MP Biomedicals Ltd., Solon, Ohio, USA). Assays for measuring CORT levels in post-DEX suppression serum, post-ACTH stimulation serum, and post-sacrifice serum were each validated by comparing parallelism between serial sample dilution curves and the assay standard curve (Sturgeon and Viljoen, 2011). The ideal sample volume was determined as the amount of sample that resulted in approximately 50% displacement of the iodinated tracer from the antibody binding sites. This volume was 0.3125 μL for post-DEX suppression serum, 0.1 μL for post-ACTH stimulation serum, and 0.2 μL for post-sacrifice serum.

### Data Analysis

Primary outcome measures were: body mass (g), plasma CORT levels (ng/ml), HPA axis responsivity (the difference between post-ACTH and post-DEX CORT levels; ng/ml); EPM open arm time (sec), entries (n), latency to enter (sec), total distance (cm), speed (cm/s), and stretch-attend bouts (n); FC freezing time (% per time bin), freezing bouts (n), motion index (arbitrary units [a.u.]). Area Under the Curve (AUC) measures were also calculated for all FC data using GraphPad Prism v10.2.3. Repeated measures ANOVAs were performed to evaluate sex (female, male) and treatment (vehicle, CBD) as between-subject factors and timepoint (acute, chronic) as the within-subject factor for HPA axis responsivity, EPM open arm time, entries and total distance, and all FC measures using JASP v0.18.3. Two-way ANOVAs were performed in JASP to evaluate plasma CORT and EPM measures and in GraphPad Prism for FC AUC measures. Mice were also classified as having high or low plasma CORT using a median split method (Henricks et al., 2019) to evaluate EPM open arm time and FC cue freezing time as a function of plasma CORT levels. Significant effects of sex were observed across multiple primary outcomes, so separate one-way ANOVAs were subsequently performed independently for each sex. Post hoc tests with Bonferroni’s correction for multiple comparisons were used. Where necessary, Greenhouse-Geisser corrections were used when Mauchly’s assumption of sphericity was violated. P < 0.05 was accepted as statistically significant. ANOVA statistics are reported in tables in the supplementary materials (Tables S1 – 4). Figures were made using GraphPad Prism and graphics were made using Inkscape v1.3. Python v3.10.9 was used to transform Ethovision positional (x,y) coordinate data into Euclidian distance from the origin point for each animal to produce the EPM trajectory maps of position over time. Euclidian distance data were then transformed into trajectory data and cumulative distributions of T(x) were graphed with these data.

## RESULTS

### CBD potentiated HPA axis physiology in female and male mice

CBD increased HPA axis responsivity compared to vehicle. This effect was apparent in both sexes, significant in males (p = 0.011), and resolved with chronic exposure (Fig 2B). HPA axis responsivity was greater after chronic exposure than acute exposure for both females and males (p’s < 0.001). Females also had greater HPA axis responsivity than males (p < 0.001). Chronic CBD did not significantly affect plasma CORT levels (Fig 2C). Females had greater plasma CORT levels than males (p = 0.001; Fig 2C inset). Daily CBD also increased body mass in females only, an effect which resolved after chronic administration (significant drug x time interaction: p < 0.001; Fig 2D-E). Average body mass for the first 12 days of measurement was significantly increased in CBD-treated female mice compared to vehicle controls (unpaired t-test: t[1,14] = 2.161, p = 0.048; Fig 2E), which was resolved by the chronic timepoint (day 14, p = 0.057; data not shown).

**Fig. 2.**
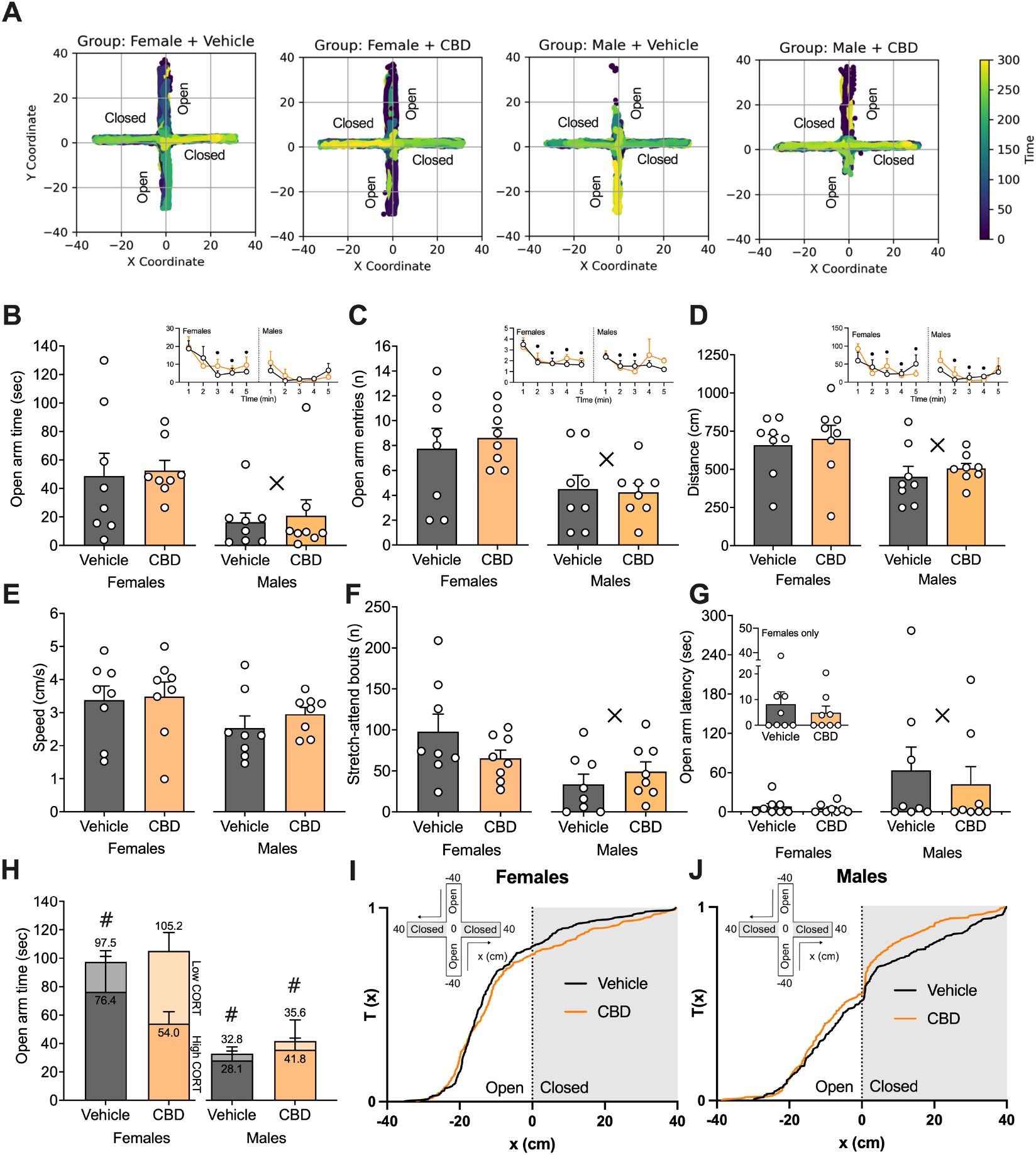
CBD effects on elevated plus maze anxiety-like behavior in female and male mice. **A)** Trajectory maps with scaled colorization for time display average position data for each group across time during EPM testing. Females explored the maze more than males and spent more time in the open arms of the maze than males. Chronic CBD did not affect **B)** open arm time, **C)** open arm entries, **D)** total distance, **E)** speed, **F)** stretch-attend posture bouts, or **G)** latency to enter the open arms. Females displayed decreased anxiety-like behaviors **(B-C**,**F-G)** and increased locomotion **(D)** compared to males. Open arm time, entries, and total distance decreased across time compared to the 1^st^ time bin (**B-D inset panels**). **H)** Mice classified using a median split as being high-CORT (solid color) had increased open arm time compared to low-CORT mice (faded color), an effect which was resolved with chronic CBD in female mice. Cumulative distribution plots for **I)** females and **J)** males map position (x) along trajectory-based axes from -40 cm (open arms) to + 40 cm (closed arms) as a function of time, T(x), to visualize sex- and drug-condition differences during EPM testing. CORT = corticosterone, CBD = cannabidiol. × = compared to female, • = compared to earlier timepoint, # = compared to low-CORT group

### CBD normalized anxiety-like behavior associated with HPA axis hypofunction in female mice

Chronic CBD did not significantly affect group averages for EPM open arm time or number of entries (Fig 2B-C) in either female or male mice (p’s > 0.05). Locomotor measures (total distance, speed) were also not affected by CBD (p’s > 0.05; Fig D-E). CBD did not affect group averages for the number of stretch-attend postures (Fig 2F) or latency to enter the open arms (Fig 2G); however, mice classified as having high plasma CORT levels had increased EPM open arm time compared to low CORT mice (p = 0.042), an effect which was normalized by chronic CBD in female, but not male, mice (Fig 2H). Significant effects of time were also observed for open arm time, entries, and total distance in females, and open arm entries and total distance in males (p’s < 0.05) indicating that maze exploration decreased with time (Fig 2A, Fig 2B-D inlets). Females had decreased anxiety-like behaviors but increased total distance compared to males (p’s < 0.05), indicating differences were locomotion-dependent. Cumulative distributions of arena position as a function of time (Fig 2I-J) demonstrate that while both male and female mice spent more time in the closed arms generally, females spent more time in the open arms than males, and chronic CBD differentially affected arm time in females vs. males.

### CBD enhanced fear responding during cued recall in female mice

Analysis of freezing time and AUCs indicated that, compared to vehicle, CBD did not significantly affect freezing behaviors during conditioning and contextual fear recall in either females or males (p’s > 0.05; Fig 3A-D). The number of freezing bouts during conditioning, context testing, and cue testing were also not significantly affected by CBD (p’s > 0.05; Fig 3A,C,E). However, significant effects of CBD were apparent after analysis of AUCs during cue testing, where CBD increased cued freezing AUC in female mice compared to vehicle (unpaired t-test: t[1,14] = 2.441, p = 0.029; Fig 3F).

**Fig. 3.**
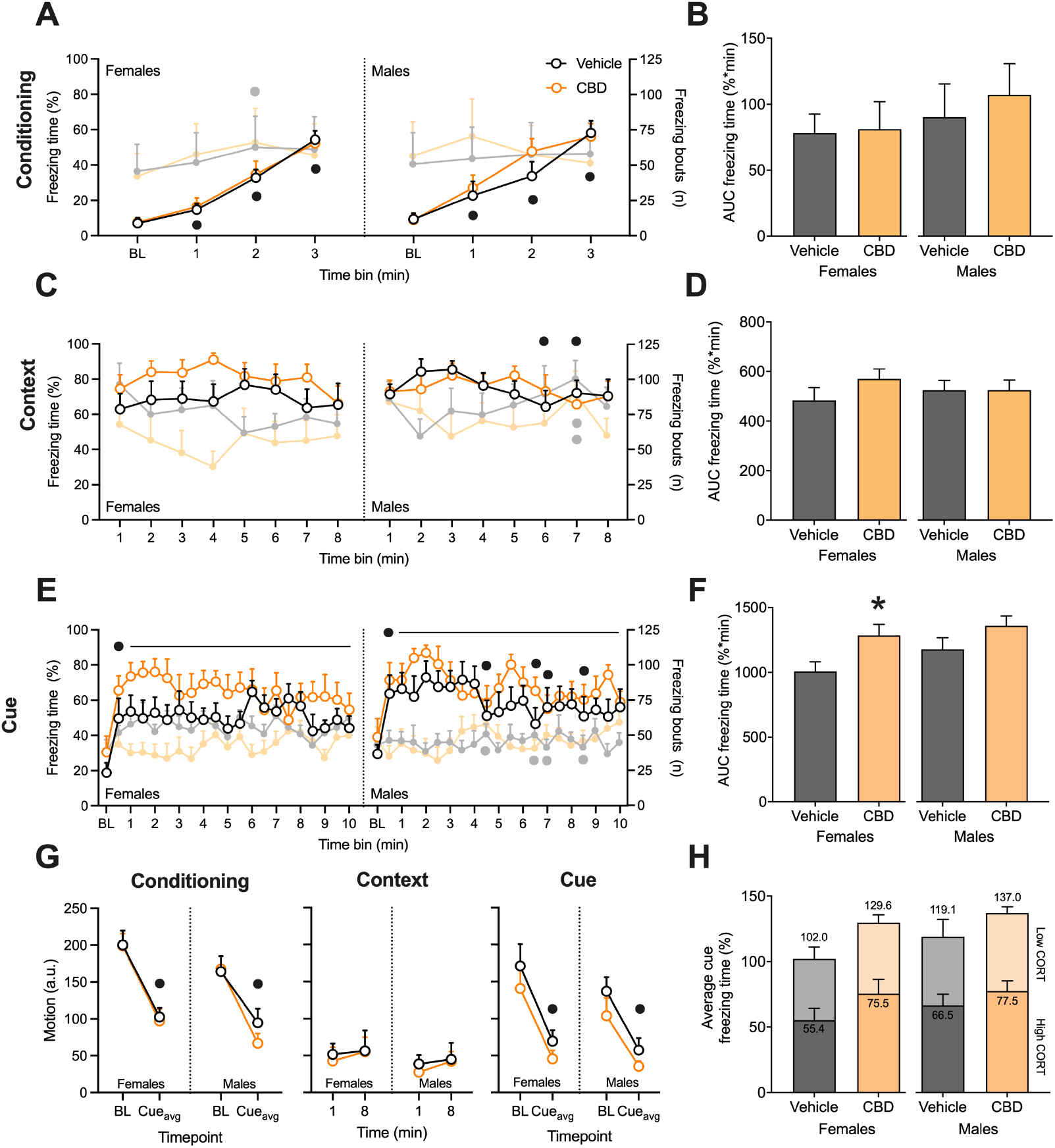
CBD effects on fear conditioning and contextual or cued fear memory recall in female and male mice. Compared to vehicle (grey time series), chronic CBD (orange time series) did not significantly affect freezing time or bouts during conditioning (**A-B**) or context testing (**C-D**). **A)** All mice increased freezing time (left y-axis, foreground time series) during conditioning; males had consistent freezing bouts (right y-axis, background faded time series) across time, while females increased freezing bouts at 2^nd^ time bin. **B)** No effect of chronic CBD on conditioning freezing time AUCs. **C)** During context testing, female**s** had consistent freezing time and bouts; male freezing time was significantly lower by the 6^th^ and 7^th^ time bin (compared to 3^rd^) and bouts were significantly greater at the 7^th^ time bin (compared to the 2^nd^ and 3^rd^). **D)** No effect of chronic CBD on context freezing time AUCs. **E)** Male and female freezing time and bouts were increased from baseline to post-cue time bins (significance represented by black horizontal lines); females maintained freezing time and bouts within-session, while male freezing time and bouts were significantly decreased at the 4.5-, 6.5-, 7-, and 8.5-minute time bins (compared to the 2^nd^ time bin). **F)** Chronic CBD significantly increased cue freezing time AUC compared to vehicle in females only. **G)** Average motion was expectedly decreased between baseline and post-cue time bins during conditioning and cue testing; motion remained low during context testing. **H)** Significant effects of chronic CBD on cue freezing time were not further dissociable by high-CORT (solid color) and low-CORT (faded color) classification. CORT = corticosterone, CBD = cannabidiol, AUC = area under the curve. * = compared to vehicle, • = compared to earlier timepoint

Sex-dependent differences in fear responding were also apparent across time. In females, main effects of time on freezing time and bouts were observed during conditioning and cue testing (p’s < 0.05) but not context testing (Fig 3A,C,E). In males, main effects of time on freezing were observed during conditioning, context testing, and cue testing (p’s < 0.01; Fig 3A,C,E). Main effects of time on freezing bouts were also observed in males during context and cue testing (p’s < 0.05; Fig 3C,E). Altogether, freezing time expectedly increased during conditioning for both females and males. Males maintained a consistent number of freezing bouts across time during conditioning (p’s > 0.05) whereas bouts were significantly greater from baseline by the second time bin in females (p = 0.021). During context testing, freezing time and bouts were consistent across time in females (p’s > 0.05). In males, freezing time decreased after the third time bin (time bins 3 vs. 6: p = 0.015; 3 vs. 7: p = 0.018) concomitant with increased freezing bouts (time bins 2 vs. 7: p = 0.038; 3 vs. 7: p = 0.035), indicating that male mice were exiting the freezing response at a greater frequency and freezing less overall within session. A significant sex x time interaction was also observed for freezing time during context testing (p = 0.03). During cue testing, freezing time and bouts were expectedly increased from baseline across every post-cue time bin in females and most post-cue time bins in males (p’s < 0.05). In males, however, freezing time and bouts decreased after the second time bin (time bins 2 vs. 4.5, 6.5, 7, 8.5: p’s < 0.05; Fig 2E) indicating short-term extinction occurred within session. Expectedly, motion during testing decreased between baseline and post-cue time bins during conditioning and cue testing (p’s < 0.001) and was lowest during context testing (p’s > 0.05; Fig 3G). Unlike anxiety-like behavior, freezing behavior during cue testing was not dissociable by high- and low-CORT classifiers (p’s > 0.05; Fig 3H).

Altogether, these results indicate that females exhibited enhanced fear responding compared to males, with sustained elevation of fear behaviors during contextual and cued fear memory recall tests. Interestingly, chronic CBD produced enhanced responding in female mice during the initial phase of cued fear memory recall in a novel context (Fig 3E-F).

## DISCUSSION

These data show that systemic CBD administration increased HPA axis responsivity, an effect which was significant in males and resolved with chronic administration. CBD also normalized anxiety-like behaviors in female mice exhibiting HPA axis hypofunction, and enhanced initial responding during cued fear memory recall in all female mice. Significant sex differences were observed throughout as female mice had greater plasma CORT levels, greater HPA axis responsivity, were more exploratory in the EPM, and were more responsive during contextual fear recall than males. These observed sex differences corroborate existing data in rodents demonstrating females have greater plasma CORT levels, display less EPM anxiety-like behaviors, and are more sensitive to stress during fear conditioning compared to males (Hassien et al., 2020; Knight et al., 2021; Solomon et al., 2015; Tinnikov, 1999). CBD did not significantly affect plasma CORT levels, associative fear learning, or contextual fear memory recall in our study. Taken together, we report that systemic CBD induced a state of HPA axis hyperactivity which was independent of existing sex differences in HPA axis physiology. This CBD-induced HPA axis hyperactivation was corroborated by a concomitant increase in body mass in females treated with CBD that resolved with chronic administration. Dysregulated HPA axis function in rodents has been correlated with increased body mass via increased appetitive drive leading to increased food intake and fat deposition (Izzi-Engbeaya et al., 2020; Malisch et al., 2007).

The observation reported herein of a novel CBD-induced HPA axis hyperactivation phenomenon was detected using consecutive DEX suppression and ACTH stimulation tests. Operationally, injected DEX suppresses HPA axis activation, consequently reducing circulating CORT levels over time, while injected ACTH stimulates the HPA axis to release endogenous CORT (Cole, 2006). The amount of circulating CORT produced (i.e., the difference between post-ACTH and post-DEX CORT levels) is quantified as HPA axis responsivity. Thus, systemic CBD in our study increased the amount of circulating CORT released after ACTH stimulation, such that we observed a greater HPA response magnitude in our CBD-treated mice at both timepoints. Our data demonstrating that plasma CORT levels were unaffected by chronic CBD is consistent with past observations that CBD does not modulate HPA-axis related gene expression under baseline (i.e., non-stressed) conditions in mice (Viudez-Martínez et al., 2018). Together, this evidence suggests that CBD potentiates the HPA axis system in stress- or experimental-activation conditions but not in non-stressed conditions. This is similarly reflected in behavioral studies reporting CBD did not alter foot-shock fear responses in unconditioned animals (Lemos et al., 2010; Montoya et al., 2020; Resstel et al., 2006; Uhernik et al., 2018).

The observed CBD-induced HPA axis hyperactivation phenomenon is supported by evidence that CBD can both directly and indirectly modulate endogenous glucocorticoid signaling. Early receptor binding assay studies in rats demonstrated that CBD displaced binding of radiolabeled DEX from the CORT binding site (Charles Eldridge & Landfield, 1990; Eldridge et al., 1991). Although doses used in these binding assay studies were not clinically relevant, these data evidence that CBD can directly influence glucocorticoid receptor occupancy and the autoregulation of HPA axis function. This explains the earlier observation that acute systemic CBD administration dose-dependently increased plasma CORT levels in experimentally naïve rats (A. Zuardi et al., 1984). CBD also indirectly influences endogenous glucocorticoid signaling mechanisms. CBD increased stress-induced HPA axis-related gene expression in mice via a 5-HT_1A_-dependent mechanism (Viudez-Martínez et al., 2018). CBD activation of HPA axis function also explains evidence that CBD normalized plasma CORT levels increased by ethanol exposure/withdrawal or delta-9-tetrahydrocannabinol (THC) exposure in adolescent and adult rats (Brancato et al., 2021, 2022; Devuono et al., 2022; Tringali et al., 2023). Together, evidence indicates that CBD can potentiate HPA axis functioning and normalize stress-activated glucocorticoid signaling.

CBD effects on HPA axis functioning also involve interactions with the endocannabinoid (eCB) system. CBD normalized eCB system and HPA axis gene expression in the nucleus accumbens and decreased EPM anxiety-like behavior in male mice experiencing heroin withdrawal (Navarrete et al., 2022). CBD is also known to inhibit eCB degradation enzyme fatty acid amide hydrolase (FAAH; (Bisogno et al., 2001) which can normalize stress-induced CORT release (McLaughlin et al., 2016). CBD regulation of FAAH activity thus may contribute to the normalization of stress-induced plasma CORT levels reported in numerous studies.

Interestingly, we also observed that chronic CBD normalized anxiety-like behaviors specifically in female mice classified as having HPA axis hypofunction. Altogether, CBD potentiates HPA axis function and modifies glucocorticoid signaling through multiple pathways to produce altered CORT levels and normalize behaviors. Future studies should characterize CBD dose-response curves and temporal dynamics for multiple biochemical, physiological, and behavioral markers of HPA axis function in both female and male animals.

The observation that CBD did not significantly affect EPM anxiety-like behavior for the other mice in our study contributes to the conflicted preclinical literature regarding the anxiolytic effects of CBD. Several preclinical studies have demonstrated that across multiple doses (0.5-50 mg/kg, i.p.), CBD decreased EPM anxiety-like behaviors (Guimarães et al., 1990; Schiavon et al., 2016; Zieba et al., 2019). In contrast, multiple doses of CBD (0-96 mg/kg, i.p.) were also previously demonstrated to have no effect on EPM anxiety-like behaviors in either female or male mice and after either acute or chronic administration (Chesworth et al., 2022; Liu et al., 2022). Discrepant results may be attributed to differences in study parameters including rodent species/strain. Interpreting EPM activity as a proxy measure for anxiety remains cautionary without careful calibration and standardization of testing methods across studies (Shoji & Miyakawa, 2021). The EPM may be a less reliable measure of anxiety-like behavior than other tests, especially when evaluating potential sex differences (Börchers et al., 2022). Given the conflicted literature, future investigations of CBD anxiolysis should include multiple assays of anxiety-like behavior to properly infer about the anxious state of the model animal and potential treatment effects.

The CBD-induced HPA axis hyperactivation we observed in this study also coincided with enhanced responding during cued fear memory recall specifically in female mice. This result contrasts existing reports that CBD decreased associative fear learning and memory in female mice (Montoya et al., 2020). However, these results are consistent with other evidence that CBD increased fear responding, albeit in male mice (Uhernik et al., 2018). Increased cued recall after CBD treatment may appear cursorily as increased fear responding and worsened outcomes. However, in the context of our reported CBD-induced HPA axis hyperactivation and effects on anxiety, enhanced fear responding during cued recall in a novel context could indicate an increased “stress elasticity” in CBD-treated females; that is, that CBD sensitized the stress response and primed animals to respond to novel stressors. In support of this, both Montoya et al., (2020) and Uhernik et al., (2018) reported increased freezing in CBD-treated male and female mice when testing cued fear memory recall in a novel context. In the latter study, the authors concluded that CBD disrupted the specific associative fear memory formation such that it potentiated stress responses to a novel context. This again supports our theory of increased stress elasticity as an emergent property of CBD potentiating HPA axis functioning. Our observation of no effect after chronic CBD during acquisition or contextual recall corroborates existing evidence with conditioning or context testing after chronic CBD administration (Chesworth et al., 2022). The extent of conditioning or the timing of CBD treatment in relation to conditioning can also both influence CBD effects on associative fear learning and memory (Song et al., 2016; Stern et al., 2017). Interestingly, CBD decreases freezing during contextual fear memory recall and extinction when the conditioning is stronger and increases freezing when conditioning is weaker (Song et al., 2016). A weaker conditioning may have similarly influenced the significant within-session cued fear extinction and CBD-enhanced cued fear responding that we observed in our study.

This study was exploratory in its design and is not without limitations that may be improved on in future studies. As mentioned above, future studies should employ multiple CBD doses to evaluate dose-related effects. Our primary aim for this exploratory study was to evaluate chronic effects of repeated exposure to a standardized dose of CBD, which is why only a single dose was selected. Future studies would also be strengthened by using a within-subject design to evaluate HPA axis responsivity and behavioral changes from baseline, as well as after acute CBD exposure. To circumvent issues with acute behavioral testing over multiple dosing days, chronic CBD exposure could begin after acute behavioral testing is complete. Finally, our results would be strengthened using additional behavioral tests to confirm effects across multiple assays and to decrease reliance on a single behavioral readout for interpretation.

In conclusion, these data provide novel evidence of a sub-behavioral phenomenon of CBD-induced HPA axis hyperactivation which occurs independently of existing sex differences in HPA axis physiology and behavior. Future dose-characterization studies should assess multi-dimensional behavioral measures of HPA axis activation at baseline and after acute administration, where possible to avoid habituation effects with repeated measures testing (Shoji & Miyakawa, 2021; Trott et al., 2022). In humans, HPA axis dysregulation is associated with anxiety, PTSD, and other neuropsychiatric disorders (Green et al., 2010; Lähdepuro et al., 2019). Females and males are also differently sensitive to the outcomes of HPA axis dysregulation (Freidenberg et al., 2010; Meewisse et al., 2007). Continued testing of CBD and other cannabis-derived compounds for sex-specific effects on HPA axis hypoactivity may reveal new mechanistic targets for medications development.

## AUTHOR CONTRIBUTIONS

Conceptualization - BWJ, HAS, AEMN, JYK; Methodology - BWJ, HAS, JYK; Software - BWJ; Formal analysis - BWJ; Investigation - BWJ, HAS, KN; Resources - AEMN, JYK; Data Curation - BWJ, HAS; Writing - Original Draft – BWJ, HAS; Writing - Review & Editing - BWJ, HAS, AEMN; Visualization - BWJ; Supervision - AEMN, JYK; Project administration - BWJ, HAS, KN, AEMN, JYK; Funding acquisition - AEMN, JYK

## Notes

Conflicts of Interest: The authors have no relevant financial or non-financial interests to disclose.

### Competing Interest Statement

The authors have declared no competing interest.

