## Supplemental Tables for "Cannabidiol (CBD) potentiates physiological and behavioral markers of hypothalamic-pituitary-adrenal (HPA) axis responsivity in female and male mice"

| *Sex-aggregated* | | Time | | | Drug | | | Sex | | | Drug x Time | | | Sex x Time | | | Sex x Drug | | | Sex x Drug x Time | | |
|---|---|---|---|---|---|---|---|---|---|---|---|---|---|---|---|---|---|---|---|---|---|---|
| | | F | DF | P | F | DF | P | F | DF | P | F | DF | P | F | DF | P | F | DF | P | F | DF | P |
| HPA axis | HPA axis responsivity | 60.61 | 1, 28 | **<0.001** | 7.64 | 1, 28 | **0.01** | 96.87 | 1, 28 | **<0.001** | 0.33 | 1, 28 | 0.57 | 2.13 | 1, 28 | 0.16 | 0.37 | 1, 28 | 0.55 | 0.12 | 1, 28 | 0.74 |
| | Body mass | 35.55 | 4.9, 138.5 | **<0.001** | 2.71 | 1, 28 | 0.11 | 108.41 | 1, 28 | **<0.001** | 1.68 | 23, 138.5 | **0.03** | 1.65 | 23, 138.5 | **0.03** | 0.17 | 1, 28 | 0.90 | 1.84 | 23, 138.5 | **0.01** |
| EPM | Open time | 7.45 | 4, 112 | **<0.001** | 0.14 | 1, 28 | 0.82 | 8.74 | 1, 28 | **0.01** | 0.21 | 4, 112 | 0.93 | 1.10 | 4, 112 | 0.36 | 0.05 | 1, 28 | 0.82 | 1.10 | 4, 112 | 0.36 |
| | Open entries | 13.62 | 4, 112 | **<0.001** | 0.15 | 1, 28 | 0.70 | 11.39 | 1, 28 | **0.002** | 0.31 | 4, 112 | 0.91 | 2.05 | 4, 112 | 0.09 | 0.08 | 1, 28 | 0.78 | 0.24 | 4, 112 | 0.91 |
| | Distance | 12.48 | 2.7, 75.1 | **<0.001** | 0.44 | 1, 28 | 0.51 | 10.35 | 1, 28 | **0.003** | 0.15 | 2.7, 75.1 | 0.92 | 1.57 | 2.7, 75.1 | 0.21 | 0.15 | 1, 28 | 0.70 | 1.91 | 2.7, 75.1 | 0.14 |
| FC | Cond freezing | 105.72 | 3, 84 | **<0.001** | 0.22 | 1, 28 | 0.65 | 1.28 | 1, 28 | 0.27 | 1.20 | 3, 84 | 0.31 | 0.70 | 3, 84 | 0.56 | 0.12 | 1, 28 | 0.73 | 0.56 | 3, 84 | 0.64 |
| | Cond bouts | 2.69 | 1, 28 | **0.05** | 0.01 | 1, 28 | 0.92 | 0.01 | 1, 28 | 0.94 | 1.07 | 1, 28 | 0.37 | 1.71 | 1, 28 | 0.17 | 0.01 | 1, 28 | 0.94 | 0.33 | 1, 28 | 0.81 |
| | Cond motion | 197.38 | 1, 28 | **<0.001** | 0.24 | 1, 28 | 0.63 | 2.73 | 1, 28 | 0.11 | 1.85 | 1, 28 | 0.18 | 1.24 | 1, 28 | 0.28 | 0.07 | 1, 28 | 0.79 | 1.07 | 1, 28 | 0.31 |
| | Context freezing | 4.04 | 4.9, 136.4 | **0.00** | 0.69 | 1, 28 | 0.42 | 0.00 | 1, 28 | 0.97 | 0.59 | 4.9, 136.4 | 0.70 | 1.31 | 4.9, 136.4 | **0.03** | 0.66 | 1, 28 | 0.43 | 2.15 | 4.9, 136.4 | 0.07 |
| | Context bouts | 2.83 | 1, 28 | **0.01** | 1.36 | 1, 28 | 0.25 | 0.85 | 1, 28 | 0.36 | 0.93 | 1, 28 | 0.48 | 1.32 | 1, 28 | 0.24 | 0.18 | 1, 28 | 0.68 | 1.78 | 1, 28 | 0.09 |
| | Context motion | 1.22 | 1, 28 | 0.28 | 0.14 | 1, 28 | 0.71 | 0.69 | 1, 28 | 0.41 | 0.21 | 1, 28 | 0.65 | 0.01 | 1, 28 | 0.92 | 0.00 | 1, 28 | 0.96 | 0.00 | 1, 28 | 0.98 |
| | Cue freezing | 7.40 | 8.8, 246.3 | **<0.001** | 2.96 | 1, 28 | 0.10 | 0.93 | 1, 28 | 0.35 | 1.18 | 8.8, 246.3 | 0.31 | 0.78 | 8.8, 246.3 | 0.63 | 0.13 | 1, 28 | 0.73 | 0.90 | 8.8, 246.3 | 0.52 |
| | Cue bouts | 8.35 | 1, 28 | **<0.001** | 2.93 | 1, 28 | 0.10 | 0.85 | 1, 28 | 0.37 | 1.26 | 1, 28 | 0.25 | 0.79 | 1, 28 | 0.64 | 0.07 | 1, 28 | 0.80 | 1.10 | 1, 28 | 0.36 |
| | Cue motion | 83.85 | 1, 28 | **<0.001** | 2.54 | 1, 28 | 0.12 | 1.85 | 1, 28 | 0.19 | 0.22 | 1, 28 | 0.64 | 1.69 | 1, 28 | 0.20 | 0.00 | 1, 28 | 1.00 | 0.02 | 1, 28 | 0.90 |

**Table S1. Statistics from two-way Repeated Measures ANOVAs of sex-aggregated data comparing effects of CBD to vehicle in female and male mice over time.** Analyses were performed to evaluate whether a) acute or chronic CBD administration would affect HPA axis responsivity or body mass, and b) whether chronic CBD administration would affect anxiety-like behavior (open arm time, entries) or locomotion (total distance) in the EPM over time or fear-related behavior (freezing time, bouts, motion) during trace fear conditioning and recall. P < 0.05 is significant (**bold text**). CBD = cannabidiol; EPM = elevated plus maze.

| *Sex-aggregated* | | CORT split | | | Drug | | | Sex | | | Drug x CORT split | | | Sex x CORT split | | | Sex x Drug | | | Sex x Drug x CORT split | | |
|---|---|---|---|---|---|---|---|---|---|---|---|---|---|---|---|---|---|---|---|---|---|---|
| | | F | DF | P | F | DF | P | F | DF | P | F | DF | P | F | DF | P | F | DF | P | F | DF | P |
| HPA axis | Plasma CORT | - | - | - | 0.05 | 1, 27 | 0.83 | 13.06 | 1, 27 | **0.001** | - | - | - | - | - | - | 0.01 | 1, 27 | 0.93 | - | - | - |
| EPM | Open time | - | - | - | 0.15 | 1, 28 | 0.70 | 8.72 | 1, 28 | **0.01** | - | - | - | - | - | - | 0.00 | 1, 28 | 0.98 | - | - | - |
| | Open entries | - | - | - | 0.08 | 1, 28 | 0.78 | 11.43 | 1, 28 | **0.002** | - | - | - | - | - | - | 0.25 | 1, 28 | 0.62 | - | - | - |
| | Distance | - | - | - | 0.51 | 1, 28 | 0.48 | 8.71 | 1, 28 | **0.01** | - | - | - | - | - | - | 0.01 | 1, 28 | 0.92 | - | - | - |
| | Speed | - | - | - | 0.51 | 1, 28 | 0.48 | 3.49 | 1, 28 | 0.07 | - | - | - | - | - | - | 0.18 | 1, 28 | 0.67 | - | - | - |
| | Stretch-attend bouts | - | - | - | 0.33 | 1, 28 | 0.57 | 7.61 | 1, 2 | **0.01** | - | - | - | - | - | - | 2.70 | 1, 28 | 0.11 | - | - | - |
| | Open time CORT split | 8.46 | 1, 24 | **0.01** | 0.19 | 1, 24 | 0.66 | 11.31 | 1, 24 | **0.003** | 1.49 | 1, 24 | 0.23 | 0.02 | 1, 24 | 0.89 | 0.00 | 1, 24 | 0.97 | 2.34 | 1, 24 | 0.14 |
| | Open latency | - | - | - | 0.30 | 1, 28 | 0.59 | 4.25 | 1, 28 | **0.05** | - | - | - | - | - | - | 0.16 | 1, 28 | 0.69 | - | - | - |
| FC | Cue freeze CORT split | 5.83 | 1, 24 | **0.02** | 3.50 | 1, 24 | 0.07 | 1.41 | 1, 24 | 0.25 | 0.33 | 1, 24 | 0.57 | 0.16 | 1, 24 | 0.70 | 0.72 | 1, 24 | 0.41 | 0.03 | 1, 24 | 0.87 |

**Table S2. Statistics from two-way ANOVAs of sex-aggregated data comparing effects of CBD to vehicle in female and male mice.** Analyses were performed to evaluate whether chronic CBD administration would affect a) plasma CORT levels, b) anxiety-like behavior (open arm time, entries, and latency to enter, and stretch-attend bouts) or locomotion (total distance, speed) measures in the EPM averaged across time, or c) whether behavioral effects of CBD are dissociable by individual differences in plasma CORT levels. P < 0.05 is significant (**bold text**). CBD = cannabidiol; EPM = elevated plus maze.

| *Sex-disaggregated* | | *Males only* Time | | | Drug | | | Drug x Time | | | *Females only* Time | | | Drug | | | Drug x Time | | |
|---|---|---|---|---|---|---|---|---|---|---|---|---|---|---|---|---|---|---|---|
| | | F | DF | P | F | DF | P | F | DF | P | F | DF | P | F | DF | P | F | DF | P |
| HPA axis | HPA axis responsivity | 76.05 | 1, 14 | **<0.001** | 8.66 | 1, 14 | **0.01** | 0.10 | 1, 14 | 0.75 | 24.61 | 1, 14 | **<0.001** | 3.28 | 1, 14 | 0.09 | 0.24 | 1, 14 | 0.63 |
| | Body mass | 17.07 | 23, 322 | **<0.001** | 0.98 | 1, 14 | 0.34 | 0.65 | 23, 322 | 0.89 | 20.63 | 23, 0 | **<0.001** | 2.94 | 1, 14 | 0.11 | 3.33 | 23, 0 | **<0.001** |
| EPM | Open time | 3.28 | 1.9, 25.9 | 0.06 | 0.04 | 1, 14 | 0.85 | 0.90 | 1.9, 25.9 | 0.41 | 4.75 | 0.0 | **0.002** | 0.14 | 0.0 | 0.71 | 0.53 | 4, 56 | 0.71 |
| | Open entries | 4.23 | 4, 56 | **0.01** | 0.32 | 1, 14 | 0.58 | 0.23 | 4, 56 | 0.92 | 10.20 | 4, 56 | **<0.001** | 0.01 | 1, 14 | 0.95 | 0.47 | 4, 56 | 0.76 |
| | Distance | 5.51 | 1.9, 25.9 | **0.01** | 0.04 | 1, 14 | 0.84 | 1.28 | 1.9, 25.9 | 0.29 | 7.58 | 4, 56 | **<0.001** | 0.49 | 1, 14 | 0.50 | 0.93 | 4, 56 | 0.45 |
| FC | Cond freezing | 40.10 | 3, 42 | **<0.001** | 0.27 | 1, 14 | 0.61 | 1.24 | 3, 42 | 0.31 | 79.60 | 3, 42 | **<0.001** | 0.01 | 1, 14 | 0.93 | 0.16 | 3, 42 | 0.92 |
| | Cond bouts | 0.81 | 2.1, 28.9 | 0.46 | 0.02 | 1, 14 | 0.90 | 1.16 | 2.1, 28.9 | 0.33 | 3.45 | 2.2, 30.8 | **0.04** | 0.00 | 1, 14 | 0.99 | 0.29 | 2.2, 30.8 | 0.77 |
| | Cond motion | 70.64 | 1, 14 | **<0.001** | 0.27 | 1, 14 | 0.61 | 2.42 | 1, 14 | 0.14 | 140.96 | 1, 14 | **<0.001** | 0.00 | 1, 14 | 0.88 | 0.07 | 1, 14 | 0.80 |
| | Context freezing | 3.30 | 7, 98 | **0.003** | 0.00 | 1, 14 | 0.99 | 1.39 | 7, 98 | 0.22 | 2.17 | 3.6, 50.1 | 0.09 | 1.12 | 1, 14 | 0.31 | 1.36 | 3.6, 50.1 | 0.26 |
| | Context bouts | 2.74 | 7, 98 | **0.01** | 0.27 | 1, 14 | 0.62 | 1.34 | 7, 98 | 0.24 | 1.47 | 7, 98 | 0.19 | 1.32 | 1, 14 | 0.27 | 1.37 | 7, 98 | 0.23 |
| | Context motion | 1.03 | 1, 14 | 0.33 | 0.14 | 1, 14 | 0.71 | 0.17 | 1, 14 | 0.69 | 0.39 | 1, 14 | 0.54 | 0.04 | 1, 14 | 0.85 | 0.07 | 1, 14 | 0.79 |
| | Cue freezing | 4.40 | 20, 280 | **<0.001** | 0.93 | 1, 14 | 0.35 | 0.91 | 20, 280 | 0.57 | 3.75 | 20, 280 | **<0.001** | 2.16 | 1, 14 | 0.16 | 1.18 | 20, 280 | 0.27 |
| | Cue bouts | 4.94 | 20, 280 | **<0.001** | 1.09 | 1, 14 | 0.31 | 0.90 | 20, 280 | 0.59 | 4.16 | 20, 280 | **<0.001** | 1.89 | 1, 14 | 0.19 | 1.50 | 20, 280 | 0.08 |
| | Cue motion | 39.18 | 1, 14 | **<0.001** | 1.55 | 1, 14 | 0.23 | 0.23 | 1, 14 | 0.64 | 45.10 | 1, 14 | **<0.001** | 1.07 | 1, 14 | 0.32 | 0.05 | 1, 14 | 0.83 |

**Table S3. Statistics from one-way Repeated Measures ANOVAs of sex-disaggregated data compared effects of CBD to vehicle separately for female and male mice.** Analyses were performed to evaluate in each sex independently whether a) acute or chronic CBD administration would affect HPA axis responsivity or body mass, and b) whether chronic CBD administration would affect anxiety-like behavior (open arm time, entries) or locomotion (total distance) in the EPM over time or fear-related behavior (freezing time, bouts, motion) during trace fear conditioning and recall. P < 0.05 is significant (**bold text**). CBD = cannabidiol; EPM = elevated plus maze.

| *Sex-disaggregated* | | *Males only* CORT split | | | Drug | | | Drug x CORT split | | | *Females only* CORT split | | | Drug | | | Drug x CORT split | | |
|---|---|---|---|---|---|---|---|---|---|---|---|---|---|---|---|---|---|---|---|
| | | F | DF | P | F | DF | P | F | DF | P | F | DF | P | F | DF | P | F | DF | P |
| HPA axis | Plasma CORT | - | - | - | 0.06 | 1, 14 | 0.82 | - | - | - | - | - | - | 0.03 | 1, 13 | 0.88 | - | - | - |
| EPM | Open time | - | - | - | 0.12 | 1, 14 | 0.73 | - | - | - | - | - | - | 0.05 | 1, 14 | 0.83 | - | - | - |
| | Open entries | - | - | - | 0.04 | 1, 14 | 0.85 | - | - | - | - | - | - | 0.23 | 1, 14 | 0.64 | - | - | - |
| | Distance | - | - | - | 0.52 | 1, 14 | 0.48 | - | - | - | - | - | - | 0.14 | 1, 14 | 0.72 | - | - | - |
| | Speed | - | - | - | 1.03 | 1, 14 | 0.33 | - | - | - | - | - | - | 0.03 | 1, 14 | 0.86 | - | - | - |
| | Stretch-attend bouts | - | - | - | 0.83 | 1, 14 | 0.38 | - | - | - | - | - | - | 1.88 | 1, 14 | 0.19 | - | - | - |
| | Open time CORT split | 5.18 | 1, 12 | **0.04** | 0.15 | 1, 12 | 0.70 | 0.06 | 1, 12 | 0.80 | 3.69 | 1, 12 | 0.08 | 0.07 | 1, 12 | 0.80 | 3.00 | 1, 12 | 0.11 |
| | Open latency | - | - | - | 0.23 | 1, 14 | 0.64 | - | - | - | - | - | - | 0.39 | 1, 14 | 0.54 | - | - | - |
| FC | Cue freeze CORT split | 3.67 | 1, 12 | 0.08 | 0.49 | 1, 12 | 0.50 | 0.25 | 1, 12 | 0.62 | 2.21 | 1, 12 | 0.16 | 3.99 | 1, 12 | 0.07 | 0.09 | 1, 12 | 0.77 |

**Table S4. Statistics from one-way ANOVAs of sex-disaggregated data comparing effects of CBD to vehicle separately for female and male mice.** Analyses were performed to evaluate in each sex independently whether chronic CBD administration would affect a) plasma CORT levels, b) anxiety-like behavior (open arm time, entries, and latency to enter, and stretch-attend bouts) or locomotion (total distance, speed) measures in the EPM averaged across time, or c) whether behavioral effects of CBD are dissociable by individual differences in plasma CORT levels. P < 0.05 is significant (**bold text**). CBD = cannabidiol; EPM = elevated plus maze.
